# Assessing nutritional pigment content and plant health status of green and red leafy vegetables by image analysis: Catching the “red herring” of plant digital color processing

**DOI:** 10.1101/2024.07.04.602112

**Authors:** Avinash Agarwal, Filipe de Jesus Colwell, Viviana Andrea Correa Galvis, Tom Hill, Neil Boonham, Ankush Prashar

## Abstract

Estimating pigment content in leafy vegetables via digital image analysis is a reliable method for real-time assessment of plant health status and nutritive value. However, the present leaf color analysis models developed using green-leafed plants do not perform reliably while analyzing images of anthocyanin-rich leaves, often giving misleading or “red herring” trends. Hence, the present study investigates variations in different digital color features for six types of leafy vegetables with varying levels of laminar anthocyanin to identify holistic digital color analysis models that could be implemented for real-time assessment of health status as well as nutritional pigment contents of leafy vegetables irrespective of laminar anthocyanin status. For this, datasets from three digital color spaces, viz., RGB (red, green, blue), HSI (hue, saturation, intensity), and *L*a*b\** (lightness, redness-greenness, yellowness-blueness), were compared with pigment contents of *n* = 320 leaf samples, and were analyzed via linear and non-linear regression using single variables, multiple linear regression, Support Vector regression, and Random Forest regression to predict chlorophyll and anthocyanin contents. While most digital color features presented abrupt shifts between anthocyanin-rich and low-anthocyanin samples, the R digital color feature did not show any deviation due to leaf anthocyanin content and was found to have the best correlation with SPAD chlorophyll meter readings (*R^2^* = 0.83). Concomitantly, H (*R^2^* = 0.82) and *a\** (*R^2^* = 0.79) features correlated most strongly with leaf anthocyanin content. In general, prediction of pigment contents was more accurate when data from all channels within a color space was analyzed simultaneously. Further, most reliable estimates of pigment content were provided by Support Vector and Random Forest regression models (0.7 < *R^2^* < 0.85). Thus, the present findings demonstrate how digital color analysis of green as well as anthocyanin-rich leafy vegetables could be implemented for assessing plant health status and nutritional pigment content non-invasively.

## 1. Introduction

Leafy vegetables are an important source of numerous nutritional components, such as antioxidants and minerals, as well as dietary fibers that promote gut health (Gupta et al., 1989; Randhawa et al., 2015; Aramrueang et al., 2019; Li et al., 2019). Owing to the growing awareness regarding their nutritional value, leafy vegetables, popularly referred to as leafy greens, have been labelled as a “superfood”. Factors such as short growth cycles, low maintenance, and ease of processing alongside the high nutritional content per unit mass have significantly influenced the shift towards leafy greens. The transition is particularly notable in this era marked by rising food demands and the saturation of arable land area. Further, rapid advancements in crop production technologies in the form of hydroponics and LED lighting over the past decade have spearheaded the increase in cultivation of leafy greens in state-of-the-art indoor vertical farms, also known as plant factories (Kozai et al., 2022). Although an omnipresent part of human diet in all parts of the world, research on leafy vegetables was largely overshadowed by the emphasis on their counterparts such as grains, fruits, and tubers, i.e., food crops that are more filling. However, with the transition of emphasis from “food quantity” to “food quality”, there is a noticeable surge in interest towards expanding and improving the production of leafy greens (Randhawa et al., 2015).

Chlorophylls (Chl) and carotenoids (Car) are two nutritional pigments that are perpetually present in all leafy vegetables. Although both these pigments have high nutritive value owing to their antioxidant capacity and their potential to act as precursors for other bioactive compounds (Olofsson et al., 2014; Pérez-Gálvez et al., 2020; Martins et al., 2023), their role as dietary biomolecules sourced from leafy greens was not extensively explored due to various reasons. For instance, research on Chl predominantly revolved around its photosynthetic potential, with limited exploration of its impact on human health due to concerns about its bioavailability (Martins et al., 2023). In contrast, although Car pigments have been well-known for their health benefits, non-leafy crops such as carrots, bell peppers, tomatoes, and pumpkins were traditionally considered as prominent sources (Mangels et al., 1993). Additionally, growing focus on improving the nutritive value of foodstuffs has highlighted the importance of anthocyanins (Anth) as key nutritional compounds owing to their high antioxidant activity and numerous potential health benefits (Yousuf et al., 2015). Consequently, there has been a concerted effort to promote large-scale production of various Anth-rich “red” leafy greens belonging to diverse plant families, including Amaranthaceae, Brassicaceae, and Lamiaceae, in response to the rising demand for highly nutritious foods (Gioia et al., 2020). Hence, leafy vegetables have emerged as crucial sources of all three of these nutritionally essential leaf pigments in the current context.

Growing interest in large-scale cultivation of Anth-rich “red-leafed” vegetables has brought to light a new challenge for growers: monitoring the health status and nutritional quality of such crops efficiently. In the past decade, machine vision technologies such as multispectral, thermal, 3D, and hyperspectral imaging have emerged as the most reliable means of high-throughput real-time monitoring of plants (Humplík et al., 2015; Waiphara et al., 2023). Amongst the different machine-vision technologies, RGB (red, green, blue)-based multispectral imaging has been adopted most widely for monitoring plant health and physiological status owing to the strong congruence between plant health, Chl content, and leaf color (Kawashima and Nakatani, 1998; Vollmann et al., 2011; Riccardi et al., 2014; Agarwal and Dutta Gupta, 2018). Over the past decades, major developments in RGB sensors have resulted in better resolution, reduced size, lower cost, easy availability, and hassle-free application of RGB cameras, making it highly feasible for commercial implementation (Bock et al., 2020; Li et al., 2020; Kim and Chung, 2021). Hence, RGB imaging has been especially well explored for crop monitoring to study the variations in leaf color which “reflect” the physiological status of plants.

Although RGB imaging primarily records data in terms of R, G, and B values, the information can be easily translated to other three-dimensional color spaces such as HSI (hue, saturation, intensity) and Commission Internationale de l’Éclairage *L*a*b\** (lightness, redness-greenness, yellowness-blueness) (Palus, 1998). The versatility of RGB data has allowed scientists to derive numerous additional color metrics, such as G-minus-R (GMR), R+G, R+G+B, and G/R (Kawashima and Nakatani, 1998; Hu et al., 2013; Wang et al., 2013). Such metrics enhance the utilization of digitized color information for crop monitoring. Further, compatibility of digitized colorimetric features with modern analytical tools such as machine learning has made in-depth plant image analysis possible (Singh et al., 2020). Hence, studies have also been conducted by implementing various machine learning algorithms for analyzing plant RGB images to estimate Chl content (Dutta Gupta and Pattanayak, 2017; Hassanijalilian et al., 2020) and identify disease symptoms (Chowdhury et al., 2021; Abbas et al., 2021; Zamani et al., 2022).

Since currently-established digital color processing protocols for monitoring plant health have been primarily developed with focus on leaf Chl and Car as key indicators of physiological status, such studies have predominantly used green-leafed plants due to the prevalence of such plants in conventional commercial cultivation (Kawashima and Nakatani, 1998; Yadav et al., 2010; Vollmann et al., 2011; Hu et al., 2013; Riccardi et al., 2014; Rigon et al., 2016; Agarwal and Dutta Gupta, 2018; Yuan et al., 2022). Conversely, the few studies carried out with red-leafed (Anth-rich) plants have primarily focused on estimating leaf Anth content (Kim and van Iersel, 2023; Clemente et al., 2023). However, co-estimation of all three types of pigments in plants remains largely unexplored, possibly owing to misleading or “red herring” trends in digital color features in the presence of high Anth concentrations. Hence, the current study aims at investigating how leaf Anth content affects digital color features by comparing the information from six different leafy vegetables with varying levels of laminar Chl and Anth. The investigation also identifies potential colorimetric attributes that may be used for assessing the health and nutritional quality for all types of plants non-invasively, irrespective of plant species or leaf Anth-status. Further, various mathematical tools, viz., linear and non-linear regressions using single variables, multiple linear regression, as well as two machine learning methods, viz., Support Vector Regression (SVR) and Random Forest Regression (RFR), have been used to generate holistic pigment content prediction models for non-invasive plant health and nutritional quality assessment.

## 2. Materials and methods

### 2.1. Plant material

Six leafy vegetables with different levels of laminar Anth content were selected for the present study (Fig. 1), and were broadly grouped as: 1) high Anth (HA) – purple basil (*Ocimum basilicum* L. var. *purpurascens*; PB) and red pak choi (*Brassica rapa* L. ssp. *chinensis* cv. ‘Rubi F1’; RPC); 2) medium Anth (MA) – scarlet kale (*Brassica oleracea* L. var. *acephala* ‘Scarlet’; SK); and 3) low Anth (LA) – green pak choi (*Brassica rapa* L. ssp. *chinensis*; PC), arugula (*Eruca vesicaria* ssp. *sativa* Mill. cv. ‘Wasabi Rocket’; WR), and Greek basil (*Ocimum basilicum* L. var. *minimum*; GB). Leaf color ranged between dark purple and reddish-green for HA samples, green with reddish-tinged lamina and red midrib for MA samples, and different shades of green with no hint of red for LA samples.

**Fig. 1.**
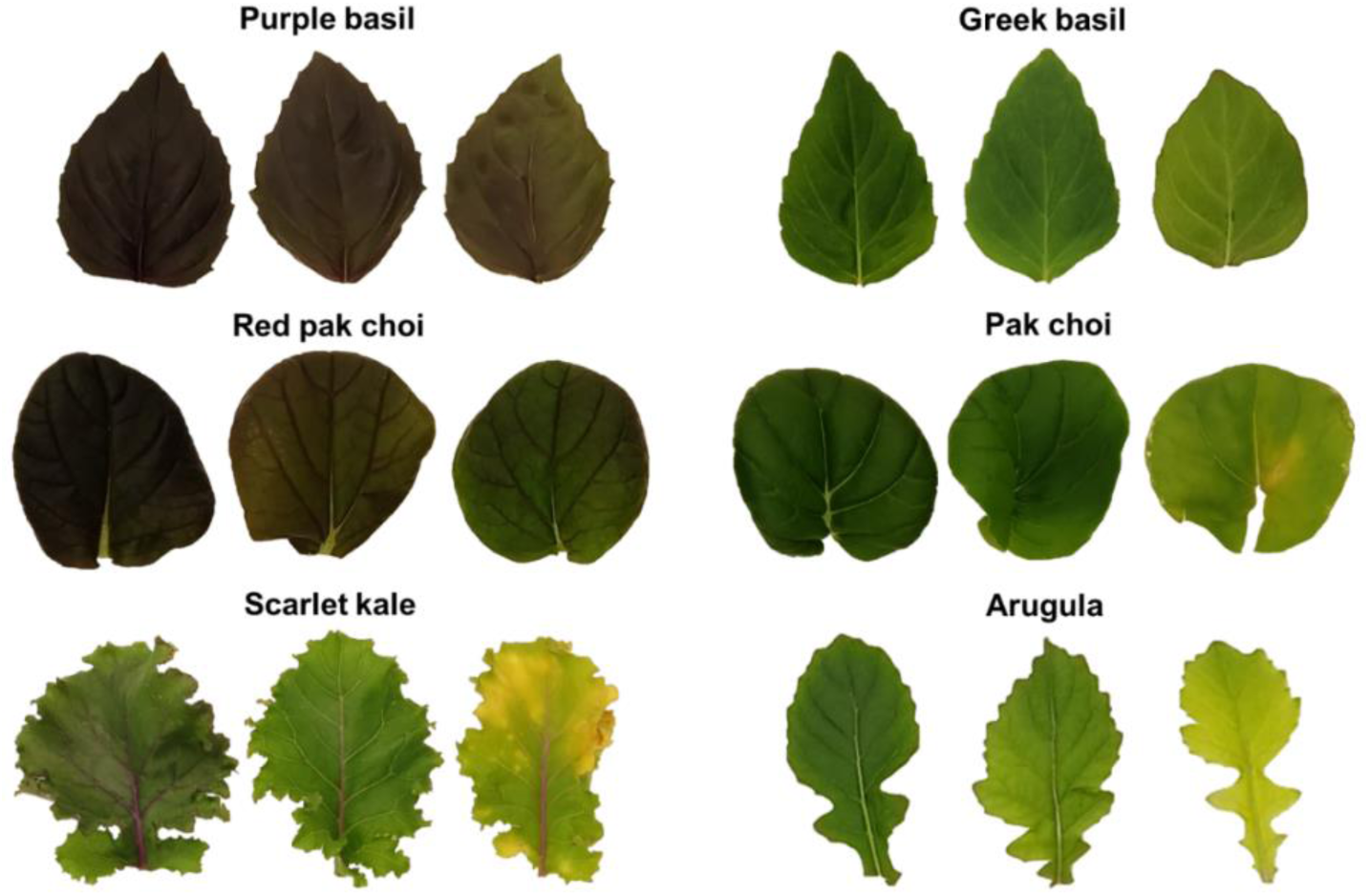
Representative leaf samples of the six types of leafy vegetables selected for the present study.

Seedlings of all plants were raised in coco-peat plugs (Van der Knapp, The Netherlands) in a nursery (Aralab-InFarm UK Ltd., London, UK), with each plug holding 5–10 seedlings. When the seedlings reached a height of *ca.* 5 cm, the seedling plugs were transferred to an experimental hydroponic vertical farm (InStore Farm V2, InFarm UK Ltd.) stationed at the Agriculture Building, Newcastle University, UK. Seedling plugs for each type of plant were placed in two hydroponic trays, where each tray (30×40 cm^2^) had a 3×4 array of equally-spaced empty slots for seedling plugs, totaling 24 seedling plugs for each plant type. A commercial hydroponics fertilizer mix was used as the nutrient source, and irrigation was performed following the ebb-and-flow system wherein the nutrient solution was flooded into the hydroponic chamber intermittently (10 min/h) to soak the roots. A white LED array having an approximate red (400–499 nm):green (500–599 nm):blue (600–699 nm) distribution of 40:20:40 was used to provide a PPFD of 280 µmol/m^2^sec following a 16/8 h day-night cycle. Temperature and relative humidity were maintained at 25±1 °C and 65±5%, respectively. Sensors for temperature, humidity, flow rate, electrical conductivity, and pH within the vertical farming system were connected to a Farmboard (InFarm UK Ltd.) for real-time monitoring of the plant growth environment.

### 2.2. Leaf sampling and SPAD measurement

Leaf samples were selected for all six types of plants (PB, *n* = 60; RPC, *n* = 40; SK, *n* = 100; PC, *n* = 40; WR, *n* = 40; GB, *n* = 40) between 15–20 days of growth within the vertical farm. Care was taken to ensure that samples with diverse levels of pigmentation were selected, whereas very young as well as fully-senesced leaves were specifically avoided. Leaves were labeled prior to excision, and three Chl meter (SPAD) readings were taken from the laminar region for each leaf using a SPAD 502 meter (Konica-Minolta, Inc., Tokyo, Japan) avoiding the midrib and prominent veins (Markwell et al., 1995). Subsequently, leaves were excised at the base for image acquisition in batches of ten, and were placed on a wet paper towel inside a sealed opaque plastic box to minimize water loss during handling and transportation, followed by sample storage for destructive measurement of pigment contents.

### 2.3. Image acquisition

Leaf samples were transferred to a customized imaging setup for digital image acquisition (Fig. 2) immediately following excision. The setup comprised of a metal frame for mounting LED-luminaires and the camera, along with a horizontal platform (stage) with a white matte surface for placing the leaves. Images were acquired using a smartphone (Redmi Note 7 Pro, Xiaomi Corp., Beijing, China) equipped with a Sony IMX 586 RGB sensor (size 1/2.0”, Quad-Bayer array) and a dual rear-camera system (primary lens: resolution 48 megapixels, aperture *f*/1.8, wide angle, pixel size 1.6 µm, phase detection autofocus; secondary lens: resolution 5 megapixels, aperture *f*/2.4, depth perception). The Open Camera android application (ver. 1.52, developer: Mark Harman, source: Google Play Store) was used for capturing images (8000×6000 pixels, sRGB color space, JPEG format). The smartphone was placed within a compact cradle suspended from the metal frame to prevent camera movement or change in camera angle. Camera-to-stage distance of 50 cm was maintained along with constant imaging parameters (exposure time 1/100 sec, ISO-200). Camera focus was fixed on the stage before placing the leaf samples, and automatic adjustments (autofocus and exposure compensation) were disabled to ensure uniformity across images. Camera flash was not used during the process to avoid glares. Instead, lighting was provided by four neutral-white (4000 K) LED tube-lights (Model No. 0051048, Feilo Sylvania International Group Kft., Budapest, Hungary; www.sylvania-lighting.com). Fixed lighting prevented undesirable fluctuations in brightness and color temperature between images, and nullified image pre-processing requirements. Since adequate lighting was provided, a relatively low ISO was adequate for ensuring sharp images while minimizing noise. Images were captured remotely using the voice-activated mode within the smartphone application to avoid shadows, delays, or any disturbances that could be caused by manual handling. During imaging, leaves were taken out of the sealed box, placed on the stage for image acquisition, and returned to the box within 1 min to ensure minimal moisture loss.

**Fig. 2.**
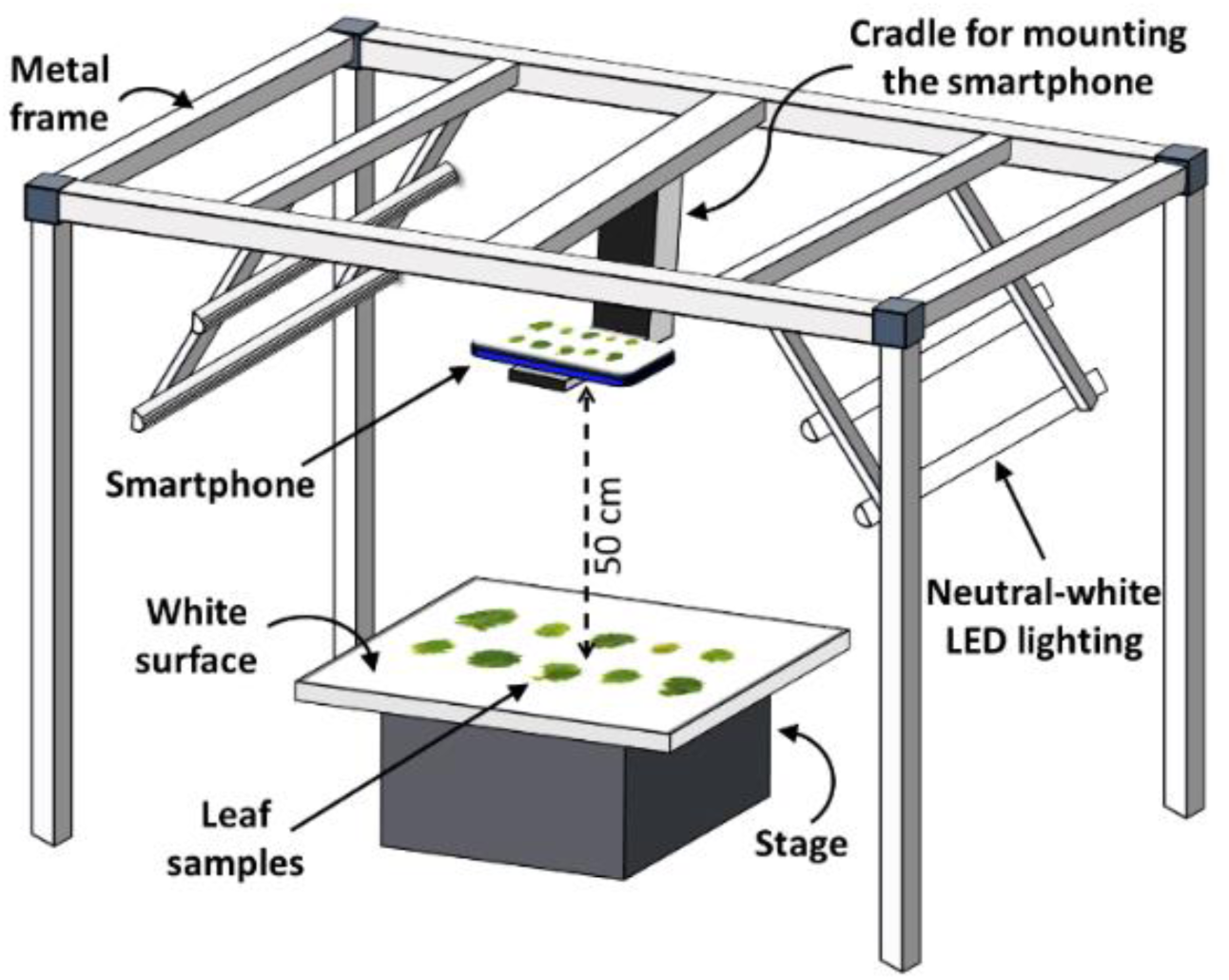
Schematic representation of the setup for leaf image acquisition.

### 2.4. Spectrophotometric estimation of pigment contents

Leaf samples were solvent-extracted for spectrophotometric estimation of total Chl, Car, and Anth contents immediately following image acquisition. Briefly, two 2 cm^2^ sections were excised from each leaf, weighed individually, sealed into separate 1.5 ml centrifuge tubes, and transferred to –20 °C for storage. Vials of all the samples were subsequently put in a liquid nitrogen bath, and pulverized with stainless-steel beads using a tissue homogenizer (Geno/Grinder 2010, SPEX SamplePrep, Cole-Parmer, Illinois, USA). One batch of vials for all six plants (*n* = 320) was used for estimating Chl and Car contents, whereas the other batch (*n* = 320) was used for estimating Anth content. Chl and Car contents was estimated following acetone extraction as described by Lichtenthaler (1987). Briefly, 1 ml of ice-cold 80% (v/v) acetone was added to each vial, followed by centrifugation at 10,000 *g* at 4 °C for 15 min. The supernatant was collected, and the pellet was re-extracted using 1 ml of the solvent. Both supernatants were pooled, and absorbance was recorded spectrophotometrically at 470 nm (A_470_), 647 nm (A_647_), and 663 nm (A_663_) for calculating the Chl and Car contents per unit leaf fresh weight (FW) per unit volume (V) of extract as follows:

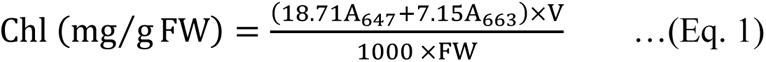

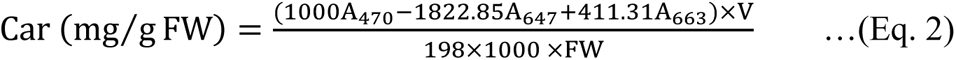

Anth content was estimated following the method of Mancinelli and Rabino (1984). A similar extraction procedure as above was followed, with chilled acidified (1% w/v HCl) methanol as the solvent. Absorbance was recorded at 530 nm (A_530_) and 657 nm (A_657_) for calculating the total Anth content using the expression A_530_ – (0.25×A_657_), where A_530_ corresponds to the peak absorbance of Anth and A_657_ was used for pheophytin correction. A standard curve was prepared using cyanidin-3-*O*-glucoside (Merck KGaA, Darmstadt, Germany) for calculating Anth content per unit leaf FW.

### 2.5. Color feature extraction

A customized image processing pipeline was designed to extract the color feature values of whole leaves using the *numpy* and *cv2* libraries in Python program (www.python.org). Within the pipeline, each RGB image was used to directly extract the R, G, B, H, S, *L\**, *a\**, and *b\** channels using standard commands, and the “I” channel values were obtained as the average of R, G, and B channels. For each channel, values for all pixels within the leaf boundary were averaged to obtain the mean color feature values for individual leaves. Three additional color indices were derived using the R and G features as follows: G/R, GMR (Wang et al., 2013), and augmented green-red index (AGRI; [GMR]×[G/R]) (Agarwal et al., 2024).

### 2.6. Data visualization and comparison

Averaged SPAD values (*n* = 3 per leaf) were plotted against the spectrophotometrically evaluated Chl content to identify global trends upon combining the data for all six plant types. Subsequently, RGB, HSI, and *L*a*b\** values were individually plotted against the SPAD values (representing Chl content) as well as the Anth content to visualize the variations in different digital color features for different levels of pigmentation. Here, SPAD values were used for comparison instead of Chl content as the former has been considered as the “gold standard” for non-invasive estimation of leaf Chl content following numerous reports over more than two decades (Wang et al., 2023). Further, the ratio of Anth and Chl contents was compared with G/R, GMR, and AGRI to understand the impact of different pigment blends on relative G and R values. For all plots, samples were labelled as HA, MA, and LA to depict the deviations induced by overall Anth content more distinctly. Curve-fitting via linear, quadratic, exponential, logarithmic, and power functions was used for global regression trends between variable pairs with all samples combined (*n* = 320). Equations along with the respective coefficient of determination (*R^2^*) for trendlines with the best fit (threshold: *R^2^* > 0.7) were presented.

### 2.7. Prediction of pigment contents by digital color features

Features from all color spaces were used for predicting pigment contents following three different approaches: 1) linear and non-linear (quadratic, logarithmic) regression using individual color features; 2) multiple linear regression using the three color spaces individually and in combination; and 3) machine learning by implementing SVR and RFR algorithms using the color spaces individually and in combination; briefly, SVR is a machine learning algorithm capable of creating linear and non-linear equations in high-dimensional space that can be fine-tuned using different hyperparameters for improving accuracy (Cortes and Vapnik, 1995), whereas RFR implements a combination of “decision trees” that depend on the values of randomly-sampled vectors to generate a “random forest” for prediction (Breiman, 2001). Linear, non-linear, and multiple linear regressions were performed using standard functions in R program (www.r-project.org), whereas SVR and RFR were performed in Python program using the *scikit-learn* library (www.scikit-learn.org). SVR was tested with three kernels or mathematical functions, viz., linear, polynomial, and Gaussian (radial basis function, *rbf*), with limited hyperparameter tuning: penalty parameter (*C*) = [0.01, 0.1, 1, 10, 100]; *rbf*-specific neighbor-impact parameter (*γ*) = [0.001, 0.01, 0.1, 1, 10]. RFR was tested with *n* = [1, 10, 100] estimators, i.e., the number of decision trees in the random forest ensemble.

### 2.8. Statistical analysis

Homogeneity of variance between the six plant types for Chl, Anth, and Car contents was assessed using the Bartlett test. Subsequently, Kruskal-Wallis non-parametric test and Dunn’s post hoc test were performed to assess the significance of difference amongst the different leafy vegetables for the content of each pigment. Goodness of fit for all prediction models was assessed in terms of the *R^2^* value. Further, root-mean-square error (RMSE) was calculated to assess the accuracy of prediction following a conventional train-test split of 80:20 (training data: *n* = 256; test data: *n* = 64), with five-fold cross-validation using different data organization sequences to simulate variations in training and test datasets.

## 3. Results

### 3.1. Comparison of pigment contents and non-invasive measurements

Chl contents showed considerable variability across all six types of plants, and although Chl concentration was significantly higher in PB (*p* < 0.05) as compared to the MA and LA plants, overlap within the ranges was observed (Fig. 3A). Similarly, while Car content was significantly higher in PB and SK (*p* < 0.05) as compared to RPC, PC, and GB, some degree of overlap in values across the six plants was evident for this pigment as well (Fig. 3A). In contrast, Anth content varied remarkably for the different types of plants (Fig. 3A) as expected. Specifically, Anth content was highest (*p* < 0.05) in the HA plants, PB followed by RPC. Further, while Anth content was detected close to the lower range in SK, it was lower still in the three LA species, i.e., GB, WR, and PC. Comparison of Chl and Car contents revealed a strong positive linear relation upon combining the data for all six plants (*R^2^* > 0.7; *n* = 320) (Fig. 3B), although no global trend was perceptible between the Anth and Chl contents due to very low Anth in MA and LA samples (Fig. 3C). Further, a strong positive correlation (*R^2^* > 0.8; *n* = 320) was observed between the Chl content and SPAD readings of the six leafy vegetables combined (Fig. 3D).

**Fig. 3.**
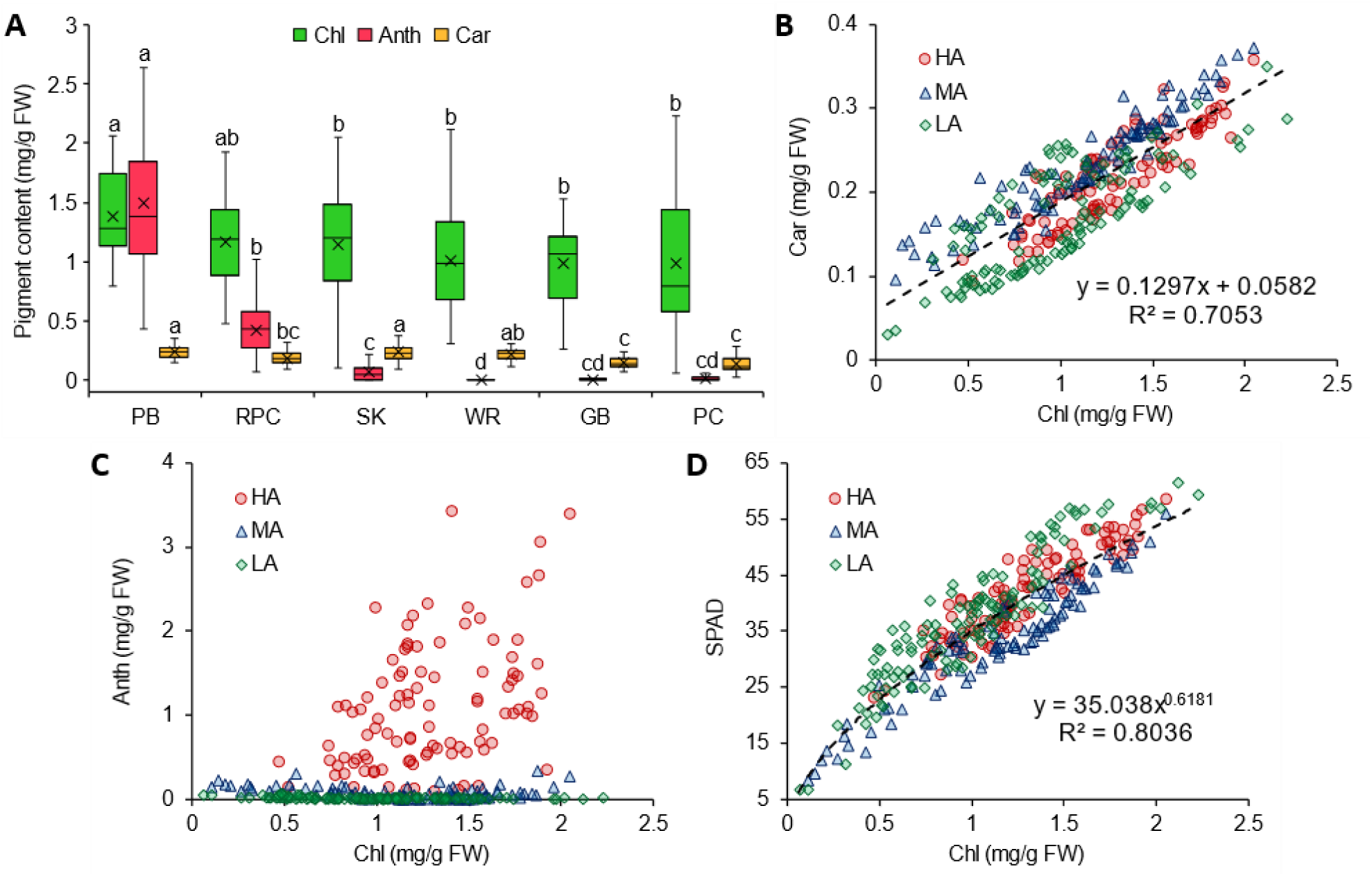
Chlorophyll (Chl), anthocyanin (Anth), and carotenoid (Car) contents of purple basil (PB; *n* = 60), red pak choi (RPC; *n* = 40), scarlet kale (SK; *n* = 100), wasabi rocket (WR; *n* = 40), Greek basil (GB; *n* = 40), and pak choi (PC; *n* = 40) (A), and the relation of Chl content with Car content (B), Anth content (C), and SPAD values (D) for plants with high (HA), medium (MA), and low (LA) levels of Anth. Box-and-Whisker plots show the mean (×), median (horizontal line), interquartile range (box), and whiskers representing 5 and 95% percentiles (A). Significant differences in mean values for each type of pigment (A) are indicated by different alphabets as per Dunn’s post hoc test (*p* < 0.05). Equations (B, D) represent trendlines for all data combined (*n* = 320). FW, fresh weight.

Plotting SPAD readings against individual color features extracted from the leaf images revealed characteristic trends with variations in pigment profile (Fig. 4). In general, HA samples clustered separately from the MA and LA samples for most of the digital color features, indicating the strong influence of leaf Anth content on digital color information. As an exception, the R color feature did not show any deviation due to the presence of Anth, and a homogenous distribution (*R^2^* = 0.83) was observed upon combining the data for all three groups of plants (Fig. 4A). Apart from R, only one other feature, viz., I, was not affected by leaf Anth content remarkably (Fig. 4F), and showed only slight deviations (*R^2^* = 0.71) compared to R. While most color features generally showed some increasing or decreasing trend with change in SPAD values, B color feature did not exhibit any distinct pattern whatsoever (Fig. 4C). Plotting the nine color features against Anth content invariably resulted in the clustering of all MA and LA samples close to the ordinate axis doe to very low total Anth content as compared to HA samples (Fig. 5). However, two color features, viz., H and *a\**, showed discernible non-linear correlations with the Anth content (Fig. 5D, H). Specifically, while H showed a strong exponential decline with increasing Anth content (*R^2^* = 0.82), *a\** exhibited a concomitant logarithmic increment (*R^2^*= 0.79). All other color features yielded relatively poor correlations (*R^2^*< 0.7) with Anth content due to abrupt transitions from LA and MA samples to the HA samples. However, color indices indicating relative changes in R and G values with changing leaf pigmentation, viz., G/R, GMR, and AGRI, exhibited distinct logarithmic decline (*R^2^* > 0.8) with increasing Anth/Chl ratio (Fig. 6).

**Fig. 4.**
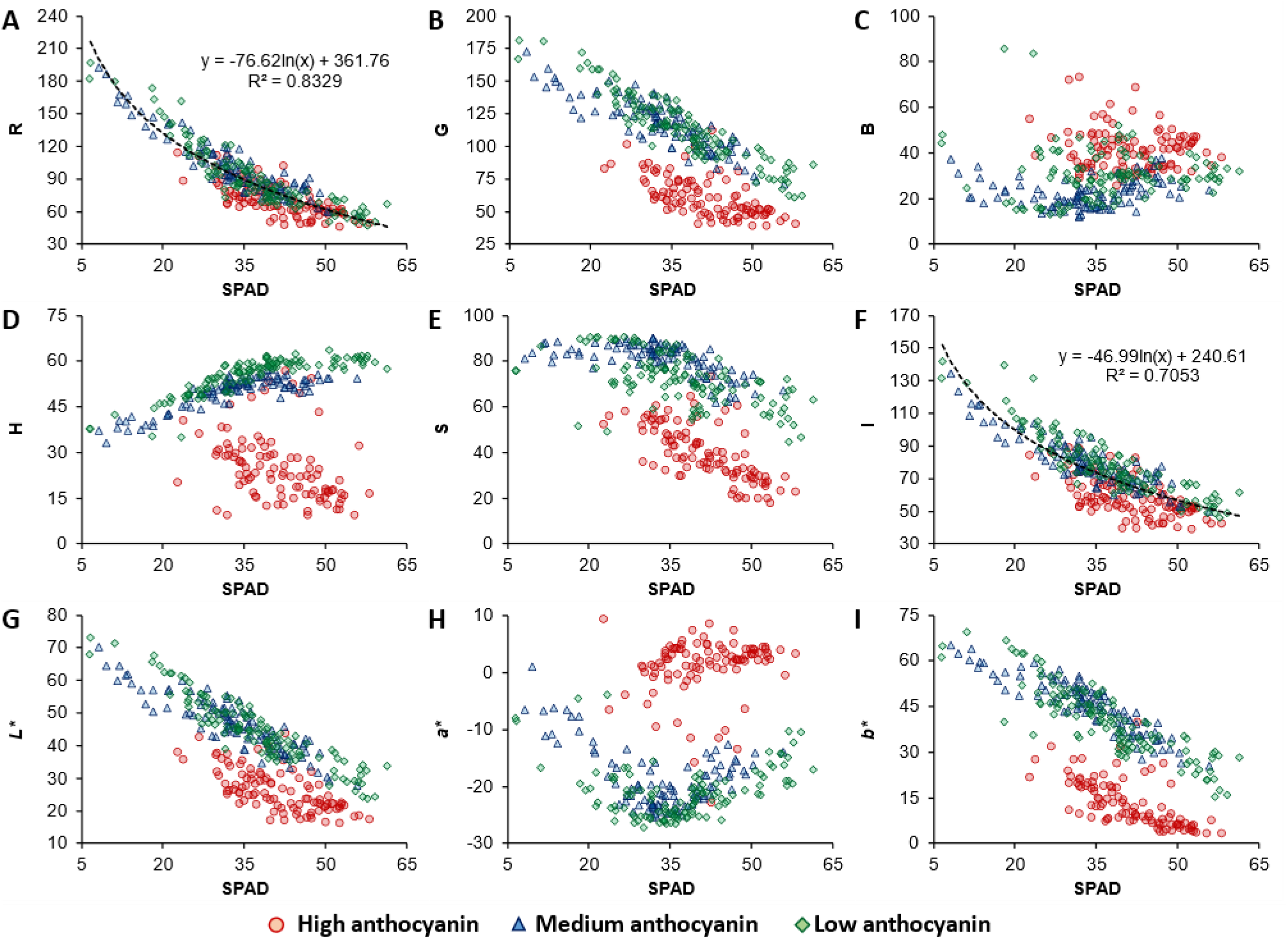
Plots of SPAD measurements with different digital color features, i.e., red (R), green (G), blue (B) (A–C), hue (H), saturation (S), intensity (I) (D–F), lightness (*L\**), redness-greenness (*a\**), and yellowness-blueness (*b\**) (G–I), for leaves with different levels of anthocyanin content. Trendlines with high coefficient of determination (*R^2^* > 0.7) have been presented, and represent samples collated from all groups (*n* = 320).

**Fig. 5.**
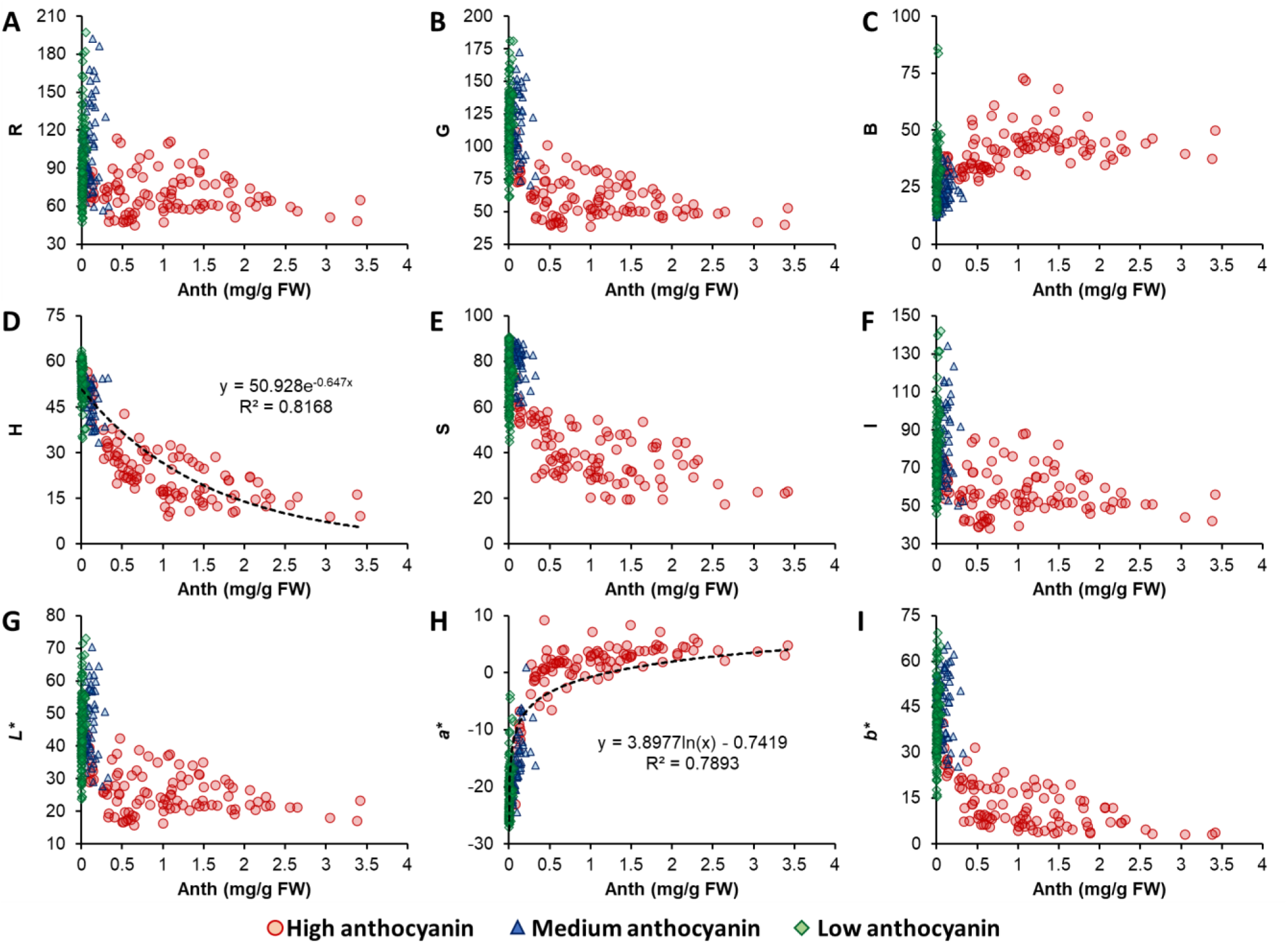
Plots of leaf anthocyanin (Anth) content with different digital color features, i.e., red (R), green (G), blue (B) (A–C), hue (H), saturation (S), intensity (I) (D–F), lightness (*L\**), redness-greenness (*a\**), and yellowness-blueness (*b\**) (G–I). Samples with different levels of Anth content have been indicated with different symbols. Trendlines with high coefficient of determination (*R^2^*> 0.7) have been presented, and represent samples collated from all groups (*n* = 320). FW, fresh weight.

**Fig. 6.**
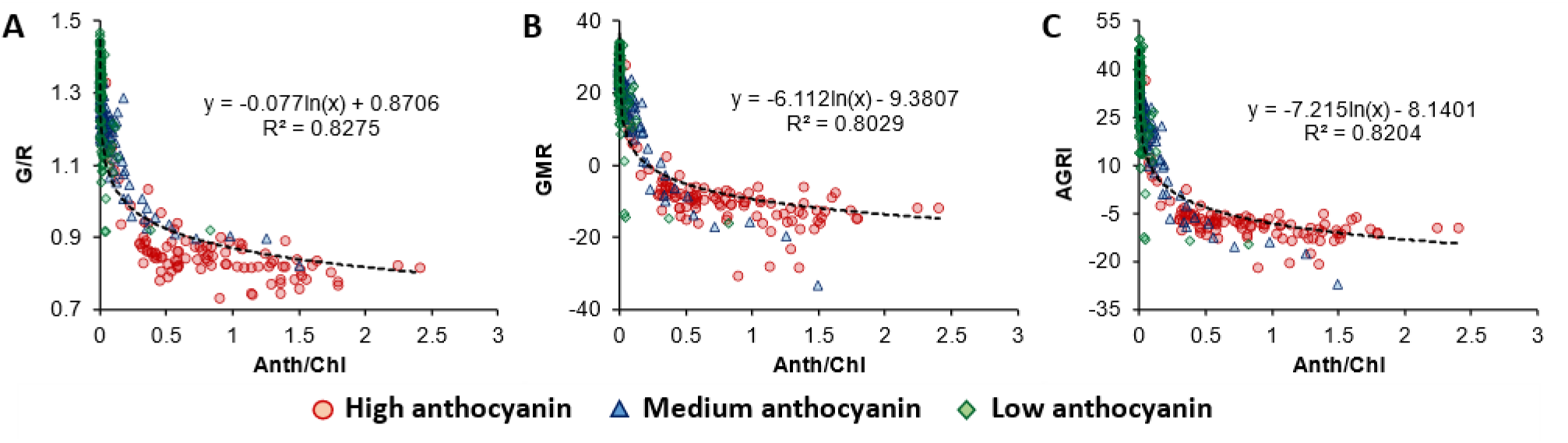
Plots of anthocyanin/chlorophyll ratio (Anth/Chl) with green/red ratio (G/R; A), green-minus-red index (GMR; B), and augmented green-red index (AGRI; C). Samples with different levels of Anth content have been indicated with different symbols. Trendlines represent samples collated from all groups (*n* = 320).

### 3.2. Predicting Chl content

Accuracy of predicting Chl content, as indicated by the *R^2^*and RMSE values, differed markedly for the different methods used, viz., linear and non-linear regression of individual color features (Table 1), multiple linear regression of trichromatic coordinates from the different color spaces (Table 2), and machine learning via SVR and RFR (Table 3). Amongst all the digital color features, estimation of Chl content was most accurate using R (*R^2^* > 0.6, RMSE < 0.27), with minor variations between the linear, quadratic, and logarithmic models (Table 1; Supplementary Table S1). This was followed by I (*R^2^* > 0.54, RMSE < 0.3), which showed better accuracy than all other features except R (Table 1). Prediction of Chl content by multiple linear regression using the trichromatic coordinates of individual color spaces yielded almost similar results (0.6 < *R^2^* < 0.62, 0.26 < RMSE < 0.27), with *L*a*b\** performing marginally better than RGB and HSI (Table 2). However, combining the information from the three color spaces resulted in considerably better prediction (*R^2^* = 0.689, RMSE = 0.243) than the individual color spaces (Table 2).

**Table 1.** Accuracy of predicting chlorophyll content using individual color features.

| Color feature | $R^2$ | | | RMSE | | |
| --- | --- | --- | --- | --- | --- | --- |
|  | *Lin* | *Quad* | *Log* | *Lin* | *Quad* | *Log* |
| **R** | 0.61 | 0.617 | 0.613 | 0.265 | 0.261 | 0.262 |
| **G** | 0.434 | 0.479 | 0.374 | 0.328 | 0.313 | 0.345 |
| **B** | 0.007 | 0.064 | 0.018 | 0.433 | 0.422 | 0.429 |
| **H** | 0.007 | 0.264 | 0.03 | 0.433 | 0.366 | 0.428 |
| **S** | 0.206 | 0.204 | 0.21 | 0.379 | 0.381 | 0.379 |
| **I** | 0.573 | 0.572 | 0.545 | 0.282 | 0.282 | 0.292 |
| **$L^*$** | 0.484 | 0.515 | 0.425 | 0.313 | 0.302 | 0.33 |
| **$a^*$** | 0.033 | 0.031 | n.a. | 0.426 | 0.427 | n.a. |
| **$b^*$** | 0.36 | 0.402 | 0.295 | 0.341 | 0.326 | 0.361 |

Accuracy parameters: $R^2$ , coefficient of determination; RMSE, root-mean-square error. Color features: R, red; G, green; B, blue; H, hue; S, saturation; I, intensity; $L^*$ , lightness; $a^*$ , redness-greenness; $b^*$ , yellowness-blueness. Models: *Lin*, linear; *Quad*, quadratic; *Log*, logarithmic. n.a., logarithmic model could not be generated due to negative values of the independent variable $a^*$ . Model equations are presented in the supplementary material.

**Table 2.** Accuracy of predicting chlorophyll (Chl) content via multiple linear regression using different color spaces.

| Color space | Equations | $R^2$ | RMSE |
| --- | --- | --- | --- |
| **RGB** | $\text{Chl} = -(0.012 \times R) + (0.0002 \times G) - (0.0005 \times B) + 2.19$ | 0.608 | 0.266 |
| **HSI** | $\text{Chl} = (0.0084 \times H) - (0.0056 \times S) - (0.015 \times I) + 2.25$ | 0.604 | 0.268 |
| **$L^*a^*b^*$** | $\text{Chl} = -(0.026 \times L^*) - (0.022 \times a^*) - (0.0087 \times b^*) + 2.14$ | 0.613 | 0.264 |
| **All_3** | $\text{Chl} = (0.297 \times R) - (0.32 \times G) - (0.06 \times B) - (0.022 \times H) - (0.0071 \times S) - (0 \times I) + (0.143 \times L^*) - (0.68 \times a^*) - (0.25 \times b^*) + 3.48$ | 0.689 | 0.243 |

Accuracy parameters: $R^2$ , coefficient of determination; RMSE, root-mean-square error. Color spaces: RGB, red, green, blue; HSI, hue, saturation, intensity; $L^*a^*b^*$ , lightness, redness-greenness, yellowness-blueness; All\_3, RGB, HSI, and $L^*a^*b^*$ combined.

**Table 3.** Accuracy of predicting chlorophyll content via machine learning (ML) using different color spaces.

| ML method | Color space | $R^2$ | | | RMSE | | |
| --- | --- | --- | --- | --- | --- | --- | --- |
| **SVR** |  | *ker_lin* | *ker_poly* | *ker_rbf* | *ker_lin* | *ker_poly* | *ker_rbf* |
|  | **RGB** | 0.602 | 0.458 | 0.694 | 0.268 | 0.311 | 0.235 |
|  | **HSI** | 0.589 | 0.587 | 0.682 | 0.272 | 0.272 | 0.24 |
|  | **$L^*a^*b^*$** | 0.606 | 0.531 | 0.652 | 0.266 | 0.291 | 0.251 |
|  | **All_3** | 0.686 | 0.701 | 0.659 | 0.238 | 0.232 | 0.248 |
| **RFR** |  | *es_1* | *es_10* | *es_100* | *es_1* | *es_10* | *es_100* |
|  | **RGB** | 0.383 | 0.635 | 0.645 | 0.332 | 0.257 | 0.253 |
|  | **HSI** | 0.501 | 0.671 | 0.702 | 0.301 | 0.244 | 0.231 |
|  | **$L^*a^*b^*$** | 0.367 | 0.639 | 0.668 | 0.338 | 0.256 | 0.245 |
|  | **All_3** | 0.454 | 0.68 | 0.692 | 0.314 | 0.24 | 0.236 |

Accuracy parameters: $R^2$ , coefficient of determination; RMSE, root-mean-square error. ML methods: SVR, support vector regression; RFR, random forest regression. Color spaces: RGB, red, green, blue; HSI, hue, saturation, intensity; $L^*a^*b^*$ , lightness, redness-greenness, yellowness-blueness; All\_3, RGB, HSI, and $L^*a^*b^*$ combined. SVR kernels: *ker\_lin*, linear; *ker\_poly*, polynomial; *ker\_rbf*, radial basis function. RFR estimators: *es\_1*, 1 estimator; *es\_10*, 10 estimators; *es\_100*, 100 estimators.

For SVR with individual colour spaces, employing the *rbf* kernel yielded higher prediction accuracy for Chl, surpassing both linear and polynomial kernels (Table 3). Specifically, the *rbf* kernel achieved an *R^2^*value exceeding 0.65, with an associated RMSE staying below 0.251. Conversely, the linear and polynomial kernels had relatively poorer performance, exhibiting *R^2^* < 0.61 and RMSE > 0.26. Herein, RGB data yielded the most favourable outcomes, showcasing an *R^2^*of 0.694 and an RMSE of 0.235. Interestingly though, amalgamating data from all three colour spaces led to a slight improvement in predictive capability, particularly noticeable when employing the polynomial kernel for regression. This integrated approach yielded an *R^2^* of 0.701 and an RMSE of 0.232, suggesting a synergistic/mutual effect resulting from the combination of multiple colour spaces. In contrast, for RFR models, predictive performance clearly improvement upon increasing the number of estimators from *n* = 1 to *n* = 10 (Table 3). Specifically, when utilizing ten estimators, *R^2^* surpassed 0.63, and the RMSE dropped below 0.26, indicating a substantial improvement. However, further increasing the number of estimators to *n* = 100 only resulted in marginal gains, with the *R^2^* exceeding 0.64 and the RMSE dropping below 0.255. In addition, data from the HSI color space resulted in the best prediction using RFR (*n* = 100 estimators: *R^2^* = 0.702, RMSE = 0.231) amongst the three color spaces, followed by the RFR model where data for all color spaces was combined (*n* = 100 estimators: *R^2^* = 0.692, RMSE = 0.236).

### 3.3. Predicting Anth content

Amongst all color features, Anth content could be predicted most accurately with H by implementing quadratic (*R^2^* = 0.786, RMSE = 0.263), logarithmic (*R^2^*= 0.768, RMSE = 0.274), and linear (*R^2^* = 0.72, RMSE = 0.301) regressions (Table 4; Supplementary Table S2). Concomitantly, quadratic regression with *a\** also resulted in reliable prediction of Anth content (*R^2^* = 0.732, RMSE = 0.294). Considering the multiple linear regression models, combining data from all color spaces provided the best prediction of Anth content (*R^2^* = 0.763, RMSE = 0.305), followed by prediction using the HSI color space data (*R^2^*= 0.739, RMSE = 0.323) (Table 5). In general, SVR models employing the *rbf* kernel provided better predictions for Anth content (*R^2^* > 0.74, RMSE < 0.32) than the linear and polynomial kernels irrespective of the color space (Table 6). Notably, predictions with HSI data yielded the best results with the *rbf* kernel (*R^2^* = 0.806, RMSE = 0.273) amongst all SVR models. However, even better predictions were obtained with RFR when the *L*a*b\** data was processed using *n* = 10 and *n* = 100 estimators (*R^2^* > 0.835, RMSE < 0.245), followed by RFR predictions with all three color spaces combined (*n* = 10 or 100 estimators: *R^2^* > 0.82, RMSE < 0.26) (Table 6).

**Table 4.** Accuracy of predicting anthocyanin content using individual color features.

| Color feature | $R^2$ | | | RMSE | | |
| --- | --- | --- | --- | --- | --- | --- |
|  | *Lin* | *Quad* | *Log* | *Lin* | *Quad* | *Log* |
| **R** | 0.097 | 0.128 | 0.116 | 0.541 | 0.531 | 0.535 |
| **G** | 0.42 | 0.533 | 0.482 | 0.433 | 0.388 | 0.409 |
| **B** | 0.29 | 0.307 | 0.28 | 0.48 | 0.473 | 0.483 |
| **H** | 0.72 | 0.786 | 0.768 | 0.301 | 0.263 | 0.274 |
| **S** | 0.577 | 0.655 | 0.627 | 0.37 | 0.334 | 0.348 |
| **I** | 0.174 | 0.229 | 0.203 | 0.517 | 0.499 | 0.508 |
| ***L*\*** | 0.359 | 0.455 | 0.411 | 0.455 | 0.42 | 0.437 |
| ***a*\*** | 0.649 | 0.732 | n.a. | 0.337 | 0.294 | n.a. |
| ***b*\*** | 0.51 | 0.676 | 0.645 | 0.398 | 0.323 | 0.339 |

Accuracy parameters: $R^2$ , coefficient of determination; RMSE, root-mean-square error. Color features: R, red; G, green; B, blue; H, hue; S, saturation; I, intensity; $L^*$ , lightness; $a^*$ , redness-greenness; $b^*$ , yellowness-blueness. Models: *Lin*, linear; *Quad*, quadratic; *Log*, logarithmic. n.a., logarithmic model could not be generated due to negative values of the independent variable $a^*$ . Model equations are presented in the supplementary material.

**Table 5.**
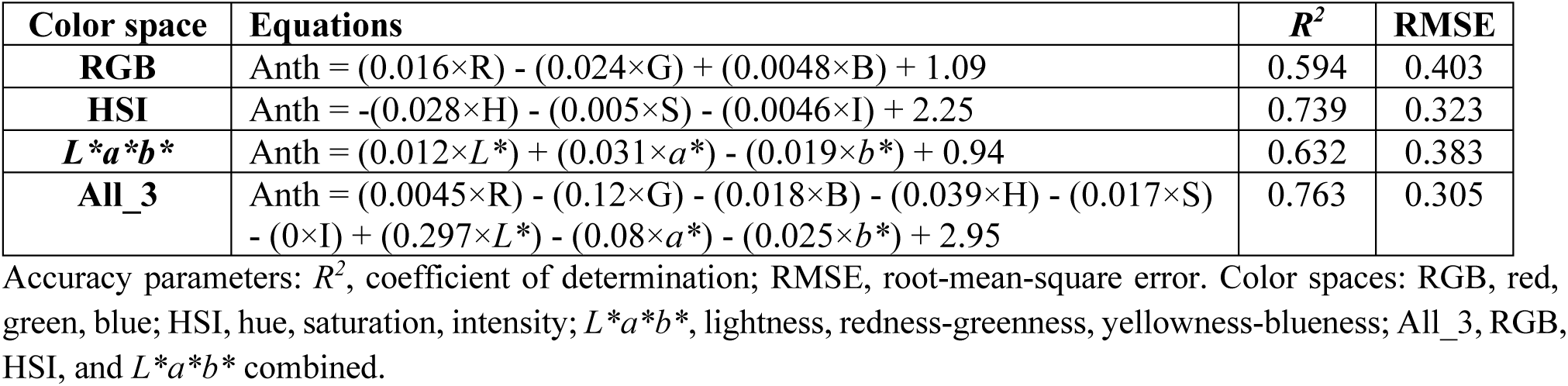
Accuracy of predicting anthocyanin (Anth) content via multiple linear regression using different color spaces.

**Table 6.** Accuracy of predicting anthocyanin content via machine learning (ML) using different color spaces.

| ML method | Color space | $R^2$ | | | RMSE | | |
| --- | --- | --- | --- | --- | --- | --- | --- |
|  |  | *ker_lin* | *ker_poly* | *ker_rbf* | *ker_lin* | *ker_poly* | *ker_rbf* |
| **SVR** | **RGB** | 0.529 | 0.448 | 0.757 | 0.433 | 0.467 | 0.307 |
|  | **HSI** | 0.713 | 0.634 | 0.806 | 0.337 | 0.381 | 0.273 |
|  | **$L^*a^*b^*$** | 0.577 | 0.383 | 0.743 | 0.41 | 0.494 | 0.317 |
|  | **All_3** | 0.712 | 0.71 | 0.761 | 0.336 | 0.339 | 0.304 |
| **RFR** |  | *es_1* | *es_10* | *es_100* | *es_1* | *es_10* | *es_100* |
|  | **RGB** | 0.605 | 0.748 | 0.755 | 0.386 | 0.314 | 0.307 |
|  | **HSI** | 0.561 | 0.758 | 0.782 | 0.407 | 0.298 | 0.284 |
|  | **$L^*a^*b^*$** | 0.649 | 0.838 | 0.847 | 0.362 | 0.243 | 0.239 |
|  | **All_3** | 0.633 | 0.824 | 0.83 | 0.378 | 0.259 | 0.252 |

Accuracy parameters: $R^2$ , coefficient of determination; RMSE, root-mean-square error. ML methods: SVR, support vector regression; RFR, random forest regression. Color spaces: RGB, red, green, blue; HSI, hue, saturation, intensity; $L^*a^*b^*$ , lightness, redness-greenness, yellowness-blueness; All\_3, RGB, HSI, and $L^*a^*b^*$ combined. SVR kernels: *ker\_lin*, linear; *ker\_poly*, polynomial; *ker\_rbf*, radial basis function. RFR estimators: *es\_1*, 1 estimator; *es\_10*, 10 estimators; *es\_100*, 100 estimators.

## 4. Discussion

Leaf pigment composition and cellular organization determines the absorption, transmission, and reflectance of incident light (Bock et al., 2020). Since leaf color is the outcome of interactions between incident visible light and the blend of pigments present, it may be considered as a dynamic attribute of the leaf which changes with physiological status. Thus, while a leaf with high Chl content appears green, leaves with very high contents of both Chl and Anth appear dark-purplish due to strong absorption of light across the entire visible spectrum (Shen et al., 2018; Hosseini et al., 2019). In contrast, a leaf devoid of all pigments appears white due to the reflection of most of the incident radiation across the entire visible waveband. In nature, this has been observed in leaves that have lost their ability to produce pigments due to genetic predispositions (Zhao et al., 2020). Since leaf pigments capture photons from specific regions of the incident light, low proportions of such photons are present in the visible radiation reflected from the leaf surface. This bestows the leaf with a specific visible color corresponding predominantly to the unabsorbed spectral region, broadly indicative of the pigments that are present at low concentrations (Sims and Gamon, 2002). Hence, the “greenness” of a healthy leaf may be attributed to the relatively low concentration of green-light-absorbing pigments such as Anth, whereas a very young leaf may appear “reddish” due to the absence of pigments primarily responsible for the absorption of red light, i.e., Chl (Gitelson and Merzlyak, 1994; Manetas, 2006; Hughes et al., 2007). Further, as all three types of dominant leaf pigments, viz., Chl, Car, and Anth, absorb blue photons, blue coloration of leaves is extremely rare.

In this context, selection of leaf samples from six different types of leafy vegetables displaying a wide range of visual color profiles (Fig. 1) enabled a comprehensive investigation of all possible deviations in digital color features due to variations in leaf pigment blends (Figs. 3–5). Most interestingly, although the “redness” of leaves is popularly associated with high Anth content, it was observed that the R color feature was not affected by Anth content at all (Fig. 4A). In contrast, G values showed a sharp reduction in the HA (Anth-rich) samples compared to the LA and MA groups (Fig. 4B). The observation can be explained by the dominant absorptive activity of Chl in the red (∼600–699 nm) and blue (∼400–499 nm) regions, and Anth in the blue-green (400–599 nm) region (Sims and Gamon, 2002). Moreover, since leaf “greenness” is commonly associated only with Chl content, a large number of digital colorimetric indices formulated for estimating Chl content utilize G (Kawashima and Nakatani, 1998; Hu et al., 2013; Wang et al., 2013; Agarwal and Dutta Gupta, 2018). Since leaf “redness” instead of “greenness” is conventionally associated with Anth content, the stark reduction in reflectance within the G channel due to increased Anth content, with no evident impact on the R channel, can be considered the “red herring” of plant digital color analysis which undermines the utility of all G-based colorimetric indices while monitoring red-leafed plants.

Analyzing the relative proportions of G and R values provided further insights into the impact of varying pigment contents on leaf color features, which would ultimately be helpful in establishing a uniform and more reliable digital color processing protocol for both red– and green-leafed plants. Increase in the relative abundance of Anth, i.e., higher Anth/Chl ratios, caused a rapid decline in G/R and GMR values until the G and R values became almost equal (*ca.* G/R = 1, GMR = 0) (Fig. 6A, B). This tendency was perceptible for AGRI as well, which is an amalgamation of G/R and GMR indices (Fig. 6C). In plants that are predominantly green-leaved, G ≤ R represents the unmasking of Car due to very low Chl content in stressed or senescing leaves (Wang et al., 2013; Agarwal et al., 2021). Since R is typically lower than G in healthy leaves due to high absorbance in the red waveband by Chl molecules (Kawashima and Nakatani, 1998; Wang et al., 2013; Dutta Gupta and Pattanayak, 2017; Widjaja Putra and Soni, 2018; Agarwal and Dutta Gupta, 2018; Agarwal et al., 2021), G/R > 1 and GMR > 0 are indicators of high Chl content and good plant health status. While all LA and a majority of MA samples showed the expected trend of G > R, interestingly, most of the HA samples had G < R, irrespective of total Chl content (Fig. 6). This deviation of relative G and R values from the established trend necessitates a reassessment of digital color indices for monitoring the health of Anth-rich plants. Based on the present data, only the R digital color feature, signifying reflectance within the red waveband, could be considered a reliable indicator of Chl content and health status for all types of plants, as it correlates strongly with Chl content while remaining totally unaffected by leaf Anth content (Fig. 4).

Considering the strong absorbance of green light by Anth, the drastic drop in G values for HA samples is understandable (Fig. 4B). However, steady reduction in G with increasing Chl content (represented using SPAD values) was prominent even in samples with lower Anth levels, viz., LA and MA (Fig. 4B). Since Chl has very low activity in the green waveband (500–599 nm), this distinct declining trend in G with increasing Chl content could be considered a misnomer. Interestingly, decline in G values with increasing Chl and Car contents has been reported in various other green-leaved plants, including wheat, rye (Kawashima and Nakatani, 1998), potato (Yadav et al., 2010), soybean (Vollmann et al., 2011; Rigon et al., 2016), rice (Hu et al., 2013), quinoa (Riccardi et al., 2014), and spinach (Agarwal et al., 2021). It may also be noted that the ratio of Chl and Car is largely conserved in healthy leaves (Gitelson, 2020). Hence, although both Chl and Car have low-level absorbance within the green waveband individually (Gitelson et al., 2002), their concomitant increase in healthy plants results in net overall reduction of green reflectance, and consequently lower G values. Additionally, in Anth-rich leaves, absorbance in the green waveband is increased further, leading to even lower G values. Hence, the ability of digital color features to indicate the content of different leaf pigments may vary according to the unique spectral attributes for each type of pigment and the blend of pigments present.

Accuracy of Chl content estimation varied markedly for the different color features (Table 1) and color space datasets (Tables 2, 3) depending on the method used for generating the regression model. As expected from the correlation trends (Fig. 4), prediction of Chl content using individual color features was most accurate with R, followed by I (Table 1). Interestingly, although the *L*a*b\** color space features did not show high predictive capacity for Chl content individually (Table 1), their prediction was better than the RGB and HSI color spaces following multiple linear regression (Table 2). This suggests that combining the information from the different channels within any color space may provide more useful information pertaining to leaf pigment contents than its individual constituent channels. Further, combining the data for the three color spaces resulted in better prediction via multiple linear regression (*R^2^*= 0.689, RMSE = 0.243). As each of the nine digital color features represents a unique aspect of leaf color, it is understandable why combining the data yielded better results via multiple linear regression than the individual color features. Notably, the coefficient for the I channel was “0” in the multiple linear regression model created by combining RGB, HSI, and *L*a*b\** datasets (Table 2). Since the values for I were calculated as the average of RGB values, the channel was likely deemed redundant by the regression algorithm in the presence of its source color features in the combined model. It is worth mentioning that preliminary tests with a fourth color space, viz., YCbCr, yielded exceptionally poor results (*R^2^* < 0.1; data not shown), and were thus not included in the present discussion.

Amongst the SVR models for each color space, Chl estimation was most accurate (*R^2^* = 0.694, RMSE = 0.235) when RGB data was processed using the *rbf* kernel (Table 3). Although the *rbf* kernel was generally more accurate than the linear and polynomial kernels for individual color spaces, combining the data from all color spaces yielded slightly better prediction via the polynomial kernel (*R^2^* = 0.701, RMSE = 0.232), followed by the linear kernel (*R^2^* = 0.686, RMSE = 0.238). This demonstrates that the efficacy of ML kernels is broadly affected by the datasets being used for training. On the other hand, accuracy of RFR models was comparable to the other prediction methods when *n* = 10 or *n* = 100 estimators were used (Table 3). However, it may be noted that while the performance of RFR models increased sharply from *n* = 1 (*R^2^*< 0.51, RMSE > 0.3) to *n* = 10 (*R^2^* > 0.63, RMSE < 0.26) estimators, it only increased slightly upon further increasing the number of estimators to *n* = 100 (*R^2^* > 0.64, RMSE < 0.255). This highlights the saturation in predictive capacity of RFR with increasing estimators. Further tests using *n* = 1000 estimators (data not shown) resulted in only slightly better accuracy than *n* = 100 estimators, albeit with higher data processing time, suggesting the pertinence of a trade-off between accuracy and processing time with increasing model estimators. Notably, RFR models with fewer estimators showed high variability in predictions for each iteration. This was because RFR is an ensemble method that utilizes a series of decision trees wherein the best subset of estimators is “randomly” chosen at each node (Liaw and Wiener, 2002). Nonetheless, higher prediction accuracy with lower variability was achieved by increasing the number of estimators. This indicates that optimizing the number of estimators in RFR not only helped in achieving higher accuracy, but also resulted in more consistent predictions.

For Anth content, prediction via quadratic regression of H (*R^2^* = 0.786, RMSE = 0.263) was found to be most accurate amongst all color features, followed by *a\** (*R^2^*= 0.732, RMSE = 0.294) (Table 4). High accuracy of H and *a\** corroborates the strong correlation of both features with Anth content (Fig. 5D, H). In contrast, prediction via all three channels in the RGB color space was relatively suboptimal (Table 4). Superior performance of H and *a\** for Anth prediction is understandable because both color features are capable of representing the transition between redness and greenness across continuous scales, whereas redness and greenness are segregated into separate channels within the RGB color space (Palus, 1998). Further, multiple linear regression predictions by the RGB data were less accurate as compared to HSI and *L*a*b\** data (Table 5), indicating that the RGB color space is not able to capture the effect of changing leaf Anth content as effectively. Hence, based on the present findings, analyzing R and G as individual color features for estimating Anth content is not recommended, although relative G versus R values can provide an insight into the relative abundance of Chl and Anth (Fig. 6). A recent study using two red-leafed lettuce cultivars demonstrated the quantification of Anth via Normalized Difference Anth Index, calculated as [I_R_ – I_G_]/[ I_R_ + I_G_], wherein I_R_ and I_G_ indicate pixel intensity in the red and green wavebands, respectively (Kim and van Iersel, 2023). A similar study by Clemente et al. (2023) elucidated Anth estimation in lettuce by using the Green Leaf Index, i.e., [2G – R – B]/[2G + R + B]. The present observations augment the findings of such studies to further elucidate more inclusive Anth prediction models that could be implemented for all types of leafy vegetables.

Compared to the quadratic regression by H (Table 4), accuracy of predicting Anth content was slightly lower for all the multiple linear regression models (*R^2^* < 0.77, RMSE > 0.3) (Table 5). However, somewhat better predictions were obtained via SVR of HSI data using the *rbf* kernel (*R^2^* = 0.806, RMSE = 0.273), whereas the most reliable prediction or Anth content amongst all models was provided by RFR of *L*a*b\** features using *n* = 10 and *n* = 100 estimators (*R^2^* > 0.835, RMSE < 0.245) (Table 6). This demonstrates that optimized pairing of regression method with color features can yield considerably more accurate Anth content predictions.

In contrast to Chl and Anth, direct estimation of Car content based on leaf reflectance is more challenging owing to the overlap of its absorbance spectra with the other leaf pigments (Gitelson et al., 2002; Sims and Gamon, 2002; Merzlyak et al., 2003), which results in extensive masking of Car. Although previous studies have utilized sophisticated instruments such as reflectance spectroscopy, mass spectrometry, and hyperspectral imaging to dissect leaf spectral traits for estimating Car content non-invasively (Merzlyak et al., 2003; Gitelson et al., 2009; Wang et al., 2019; Sonobe et al., 2020; Sousa, 2022), the intricacies of data processing presented therein might create an impediment while transferring these technologies to commercial farming operations with diverse plants. As observed in the present study, the situation was further complicated in the presence of high concentrations of Anth which resulted in non-uniform trends for almost all digital color features except R (Fig. 4).

However, estimation of Car content by using Chl content and Chl/Car ratio could be a viable option considering the high stability of Chl/Car ratio under normal physiological conditions (Gitelson, 2020; Song and Wang, 2022). In concurrence with previous studies, our findings also indicated a strong positive relation between the contents of both these pigments (*R^2^* = 0.71) upon collating the data for all six plants (Fig. 3B). Furthermore, correlating Chl and Car contents for each type of plant individually revealed even stronger correlations between both pigments (0.72 < *R^2^* < 0.93), with the Chl/Car ratio varying between *ca.* 5.5–7.5 depending upon the plant species (data not shown). Hence, considering the strong and conserved positive correlation between Chl and Car, indirect estimation of Car content by evaluating Chl content using RGB images could be deemed as the most reliable approach for assessing Car content in Anth-rich leaves, wherein knowledge of species-specific Chl/Car ratios would enable highly accurate estimates.

## 5. Conclusion

As the interest in Anth-rich leafy vegetables is growing steadily, improvement and optimization of high-throughput approaches for assessing the health and nutritional quality of such plants has become imperative. In the present study, data for multiple plants possessing varying levels of laminar Anth content was simultaneously analyzed to provide a broad-spectrum overview of variations in different digital color features commonly used for plant image analysis. Color features such as R, H, and *a\** were found to correlate strongly with leaf pigment contents, and hence, could be deployed for developing reliable colorimetric indices for real-time non-invasive health and nutritional assessment of both green– and red-leafed plants. In depth analyses with indices developed using multiple color features would provide further insights into the possibilities of digital color analysis for Anth-rich plants. As the present study was carried out under fixed lighting, image pre-processing was not needed. However, experiments under different lighting regimes would be beneficial in understanding how external light will affect the color features in such plants. Additionally, improving prediction accuracy via more elaborate and advanced data processing tools such as deep-learning will create more avenues for applying the knowledge in commercial growing systems.

## Acknowledgments

We thank the InFarm UK team for supplying seedlings and providing technical support, along with InFarm Crop Science team (Germany) for their support. We acknowledge all the partners (RoboScientific, Marks and Spencer, and InFarm) for their feedback and support in the project. We also thank the staff at Newcastle University for their technical, administrative and logistic support.

## Author contributions

Avinash Agarwal: Conceptualization, Methodology, Software, Investigation, Formal analysis, Data curation, Writing-Original draft preparation, Visualization; Filipe de Jesus Colwell: Resources, Methodology, Writing-Reviewing and Editing; Viviana Andrea Correa Galvis: Supervision, Writing-Reviewing and Editing, Project administration, Funding acquisition, Resources; Tom Hill: Supervision, Writing-Reviewing and Editing, Project administration, Funding acquisition, Resources; Neil Boonham: Supervision, Writing-Reviewing and Editing, Project administration, Funding acquisition, Resources; Ankush Prashar: Conceptualization, Supervision, Validation, Writing-Reviewing and Editing, Funding acquisition.

## References

1. Abbas, A., Jain, S., Gour, M., & Vankudothu, S. (2021). Tomato plant disease detection using transfer learning with C-GAN synthetic images. Computers and Electronics in Agriculture, 187, 106279. 10.1016/j.compag.2021.106279

2. Agarwal, A., de Jesus Colwell, F., Bello Rodriguez, J., Sommer, S., Correa Galvis, V. A., Hill, T., Boonham, N., & Prashar, A. (2024). Monitoring root rot in flat-leaf parsley via machine vision by unsupervised multivariate analysis of morphometric and spectral parameters. European Journal of Plant Pathology. 10.1007/s10658-024-02834-z

3. Agarwal, A., Dongre, P. K., & Dutta Gupta, S. (2021). Smartphone-assisted real-time estimation of chlorophyll and carotenoid concentrations and ratio using the inverse of red and green digital color features. Theoretical and Experimental Plant Physiology, 33(3), 293–302. 10.1007/s40626-021-00210-4

4. Agarwal, A., & Dutta Gupta, S. (2018). Assessment of spinach seedling health status and chlorophyll content by multivariate data analysis and multiple linear regression of leaf image features. Computers and Electronics in Agriculture, 152, 281–289. 10.1016/j.compag.2018.06.048

5. Aramrueang, N., Asavasanti, S., & Khanunthong, A. (2019). Leafy vegetables. In Integrated Processing Technologies for Food and Agricultural By-Products (pp. 245–272). Elsevier. 10.1016/B978-0-12-814138-0.00010-1

6. Bock, C. H., Barbedo, J. G. A., Del Ponte, E. M., Bohnenkamp, D., & Mahlein, A.-K. (2020). From visual estimates to fully automated sensor-based measurements of plant disease severity: status and challenges for improving accuracy. Phytopathology Research, 2, 9. 10.1186/s42483-020-00049-8

7. Breiman, L. (2001). Random Forests. Machine Learning, 45, 5–32.

8. Chowdhury, M. E. H., Rahman, T., Khandakar, A., Ayari, M. A., Khan, A. U., Khan, M. S., Al-Emadi, N., Reaz, M. B. I., Islam, M. T., & Ali, S. H. M. (2021). Automatic and reliable leaf disease detection using deep learning techniques. AgriEngineering, 3(2), 294–312. 10.3390/agriengineering3020020

9. Clemente, A. A., Maciel, G. M., Siquieroli, A. C. S., Gallis, R. B. de A., Luz, J. M. Q., Sala, F. C., Pereira, L. M., & Yada, R. Y. (2023). Nutritional characterization based on vegetation indices to detect anthocyanins, carotenoids, and chlorophylls in mini-lettuce. Agronomy, 13, 1403. 10.3390/agronomy13051403

10. Cortes, C., & Vapnik, V. (1995). Support-vector networks. Machine Learning, 20, 273–297. 10.1007/BF00994018

11. Dutta Gupta, S., & Pattanayak, A. K. (2017). Intelligent image analysis (IIA) using artificial neural network (ANN) for non-invasive estimation of chlorophyll content in micropropagated plants of potato. In Vitro Cellular and Developmental Biology – Plant, 53(6), 520–526. 10.1007/s11627-017-9825-6

12. Gioia, F. Di, Tzortzakis, N., Rouphael, Y., Kyriacou, M. C., Sampaio, S. L., Ferreira, I. C. F. R., & Petropoulos, S. A. (2020). Grown to be blue—antioxidant properties and health effects of colored vegetables. Part II: Leafy, fruit, and other vegetables. Antioxidants, 9, 97. 10.3390/antiox9020097

13. Gitelson, A. (2020). Towards a generic approach to remote non-invasive estimation of foliar carotenoid-to-chlorophyll ratio. Journal of Plant Physiology, 252, 153227. 10.1016/j.jplph.2020.153227

14. Gitelson, A. A., Chivkunova, O. B., & Merzlyak, M. N. (2009). Nondestructive estimation of anthocyanins and chlorophylls in anthocyanic leaves. American Journal of Botany, 96(10), 1861– 1868. 10.3732/ajb.0800395

15. Gitelson, A. A., Zur, Y., Chivkunova, O. B., & Merzlyak, M. N. (2002). Assessing carotenoid content in plant leaves with reflectance spectroscopy. Photochemistry and Photobiology, 75(3), 272–281.

16. Gitelson, A., & Merzlyak, M. N. (1994). Quantitative estimation of chlorophyll-a using reflectance spectra: Experiments with autumn chestnut and maple leaves. Journal of Photochemistry and Photobiology B: Biology, 22, 247–252.

17. Gupta, K., Barat, G. K., Wagle, D. S., & Chawla, H. K. L. (1989). Nutrient contents and antinutritional factors in conventional and non-conventional leafy vegetables. Food Chemistry, 31, 105–116.

18. Hassanijalilian, O., Igathinathane, C., Doetkott, C., Bajwa, S., Nowatzki, J., & Haji Esmaeili, S. A. (2020). Chlorophyll estimation in soybean leaves infield with smartphone digital imaging and machine learning. Computers and Electronics in Agriculture, 174, 105433. 10.1016/j.compag.2020.105433

19. Hosseini, A., Zare Mehrjerdi, M., Aliniaeifard, S., & Seif, M. (2019). Photosynthetic and growth responses of green and purple basil plants under different spectral compositions. Physiology and Molecular Biology of Plants, 25(3), 741–752. 10.1007/s12298-019-00647-7

20. Hu, H., Zhang, J., Sun, X., & Zhang, X. (2013). Estimation of leaf chlorophyll content of rice using image color analysis. Canadian Journal of Remote Sensing, 39(2), 185–190. 10.5589/m13-026

21. Hughes, N. M., Morley, C. B., & Smith, W. K. (2007). Coordination of anthocyanin decline and photosynthetic maturation in juvenile leaves of three deciduous tree species. New Phytologist, 175(4), 675–685. 10.1111/j.1469-8137.2007.02133.x

22. Humplík, J. F., Lazár, D., Husičková, A., & Spíchal, L. (2015). Automated phenotyping of plant shoots using imaging methods for analysis of plant stress responses – A review. Plant Methods, 11, 29. 10.1186/s13007-015-0072-8

23. Kawashima, S., & Nakatani, M. (1998). An algorithm for estimating chlorophyll content in leaves using a video camera. Annals of Botany, 81, 49–54.

24. Kim, C., & van Iersel, M. W. (2023). Image-based phenotyping to estimate anthocyanin concentrations in lettuce. Frontiers in Plant Science, 14, 1155722. 10.3389/fpls.2023.1155722

25. Kim, J. Y., & Chung, Y. S. (2021). A short review of RGB sensor applications for accessible high-throughput phenotyping. Journal of Crop Science and Biotechnology, 24(5), 495–499. 10.1007/s12892-021-00104-6

26. Kozai, T., Amagai, Y., Lu, N., Hayashi, E., Ibaraki, Y., Takagaki, M., Shinohara, Y., & Maruo, T. (2022). Toward commercial production of head vegetables in plant factories with artificial lighting. In T. Kozai, G. Niu, & J. Masabni (Eds.), Plant Factory Basics, Applications and Advances (pp. 417– 434). Academic Press. 10.1016/B978-0-323-85152-7.00019-7

27. Li, D., Li, C., Yao, Y., Li, M., & Liu, L. (2020). Modern imaging techniques in plant nutrition analysis: A review. Computers and Electronics in Agriculture, 174, 105459. 10.1016/j.compag.2020.105459

28. Li, Y., Cui, Y., Lu, F., Wang, X., Liao, X., Hu, X., & Zhang, Y. (2019). Beneficial effects of a chlorophyll-rich spinach extract supplementation on prevention of obesity and modulation of gut microbiota in high-fat diet-fed mice. Journal of Functional Foods, 60, 103436. 10.1016/j.jff.2019.103436

29. Liaw, A., & Wiener, M. (2002). Classification and Regression by randomForest. R News, 2, 18–22.

30. Lichtenthaler, H. K. (1987). Chlorophylls and carotenoids: Pigments of photosynthetic biomembranes. 350–382. 10.1016/0076-6879(87)48036-1

31. Mancinelli, A. L., & Rabino, I. (1984). Photoregulation of anthocyanin synthesis X. Dependence on photosynthesis of high irradiance response anthocyanin synthesis in *Brassica oleracea* leaf disks and *Spirodela polyrrhiza*. Plant & Cell Physiology, 25(7), 1153–1160.

32. Manetas, Y. (2006). Why some leaves are anthocyanic and why most anthocyanic leaves are red? *Flora: Morphology, Distribution*, Functional Ecology of Plants, 201(3), 163–177. 10.1016/j.flora.2005.06.010

33. Mangels, A. R., Holden, J. M., Beecher, G. R., Forman, M. R., & Lanza, E. (1993). Carotenoid content of fruits and vegetables: An evaluation of analytic data. Journal of the American Dietetic Association, 93, 284–296.

34. Markwell, J., Osterman, J. C., & Mitchell, J. L. (1995). Calibration of the Minolta SPAD-502 leaf chlorophyll meter. Photosynthesis Research, 46, 467–472.

35. Martins, T., Barros, A. N., Rosa, E., & Antunes, L. (2023). Enhancing health benefits through chlorophylls and chlorophyll-rich agro-food: A comprehensive review. Molecules, 28, 5344. 10.3390/molecules28145344

36. Merzlyak, M. N., Gitelson, A. A., Chivkunova, O. B., Solovchenko, A. E., & Pogosyan, S. I. (2003). Application of reflectance spectroscopy for analysis of higher plant pigments. Russian Journal of Plant Physiology, 50, 785–792.

37. Olofsson, P., Hultqvist, M., Hellgren, L. I., & Holmdahl, R. (2014). Phytol: A chlorophyll component with anti-inflammatory and metabolic properties. In C. Jacob, G. Kirsch, A. Slusarenko, P. G. Winyard, & T. Burkholz (Eds.), Recent Advances in Redox Active Plant and Microbial Products: From Basic Chemistry to Widespread Applications in Medicine and Agriculture (pp. 345–359). Springer Netherlands. 10.1007/978-94-017-8953-0_13

38. Palus, H. (1998). Representations of colour images in different colour spaces. In The Colour Image Processing Handbook (pp. 67–90). Springer US. 10.1007/978-1-4615-5779-1_4

39. Pérez-gálvez, A., Viera, I., & Roca, M. (2020). Carotenoids and chlorophylls as antioxidants. Antioxidants, 9, 505. 10.3390/antiox9060505

40. Randhawa, M. A., Khan, A. A., Javed, M. S., & Sajid, M. W. (2015). Green leafy vegetables: A health promoting source. In Handbook of Fertility: Nutrition, Diet, Lifestyle and Reproductive Health (pp. 205–220). Elsevier Inc. 10.1016/B978-0-12-800872-0.00018-4

41. Riccardi, M., Mele, G., Pulvento, C., Lavini, A., D’Andria, R., & Jacobsen, S. E. (2014). Non-destructive evaluation of chlorophyll content in quinoa and amaranth leaves by simple and multiple regression analysis of RGB image components. Photosynthesis Research, 120(3), 263–272. 10.1007/s11120-014-9970-2

42. Rigon, J. P. G., Capuani, S., Fernandes, D. M., & Guimarães, T. M. (2016). A novel method for the estimation of soybean chlorophyll content using a smartphone and image analysis. Photosynthetica, 54(4), 559–566. 10.1007/s11099-016-0214-x

43. Shen, J., Zou, Z., Zhang, X., Zhou, L., Wang, Y., Fang, W., & Zhu, X. (2018). Metabolic analyses reveal different mechanisms of leaf color change in two purple-leaf tea plant (Camellia sinensis L.) cultivars. Horticulture Research, 5, 7. 10.1038/s41438-017-0010-1

44. Sims, D. A., & Gamon, J. A. (2002). Relationships between leaf pigment content and spectral reflectance across a wide range of species, leaf structures and developmental stages. Remote Sensing of Environment, 81, 337–354. www.elsevier.com/locate/rse

45. Singh, V., Sharma, N., & Singh, S. (2020). A review of imaging techniques for plant disease detection. Artificial Intelligence in Agriculture, 4, 229–242. 10.1016/j.aiia.2020.10.002

46. Song, G., & Wang, Q. (2022). Developing hyperspectral indices for assessing seasonal variations in the ratio of chlorophyll to carotenoid in deciduous forests. Remote Sensing, 14, 1324. 10.3390/rs14061324

47. Sonobe, R., Yamashita, H., Mihara, H., Morita, A., & Ikka, T. (2020). Estimation of leaf chlorophyll a, b and carotenoid contents and their ratios using hyperspectral reflectance. Remote Sensing, 12, 3265. 10.3390/rs12193265

48. Sousa, C. (2022). Anthocyanins, carotenoids and chlorophylls in edible plant leaves unveiled by tandem mass spectrometry. Foods, 11, 1924. 10.3390/foods11131924

49. Vollmann, J., Walter, H., Sato, T., & Schweiger, P. (2011). Digital image analysis and chlorophyll metering for phenotyping the effects of nodulation in soybean. Computers and Electronics in Agriculture, 75, 190–195. 10.1016/j.compag.2010.11.003

50. Waiphara, P., Bourgenot, C., Compton, L. J., & Prashar, A. (2022). Optical imaging resources for crop phenotyping and stress detection. In P. Duque & D. Szakonyi (Eds.), Methods in Molecular Biology (Vol. 2494, pp. 255–265). Humana. 10.1007/978-1-0716-2297-1_18

51. Wang, Y., Hu, X., Jin, G., Hou, Z., Ning, J., & Zhang, Z. (2019). Rapid prediction of chlorophylls and carotenoids content in tea leaves under different levels of nitrogen application based on hyperspectral imaging. Journal of the Science of Food and Agriculture, 99(4), 1997–2004. 10.1002/jsfa.9399

52. Wang, Y., Wang, D., Zhang, G., & Wang, J. (2013). Estimating nitrogen status of rice using the image segmentation of G-R thresholding method. Field Crops Research, 149, 33–39. 10.1016/j.fcr.2013.04.007

53. Wang, Y., Yang, Z., Gert, K., & Khan, H. A. (2023). The impact of variable illumination on vegetation indices and evaluation of illumination correction methods on chlorophyll content estimation using UAV imagery. Plant Methods, 19, 51. 10.1186/s13007-023-01028-8

54. Widjaja Putra, B. T., & Soni, P. (2018). Enhanced broadband greenness in assessing Chlorophyll a and b, Carotenoid, and Nitrogen in Robusta coffee plantations using a digital camera. Precision Agriculture, 19(2), 238–256. 10.1007/s11119-017-9513-x

55. Yadav, S. P., Ibaraki, Y., & Dutta Gupta, S. (2010). Estimation of the chlorophyll content of micropropagated potato plants using RGB based image analysis. Plant Cell, Tissue and Organ Culture, 100(2), 183–188. 10.1007/s11240-009-9635-6

56. Yousuf, B., Gul, K., Wani, A. A., & Singh, P. (2016). Health benefits of anthocyanins and their encapsulation for potential use in food systems: A review. Critical Reviews in Food Science and Nutrition, 56(13), 2223–2230. 10.1080/10408398.2013.805316

57. Yuan, Y., Wang, X., Shi, M., & Wang, P. (2022). Performance comparison of RGB and multispectral vegetation indices based on machine learning for estimating Hopea hainanensis SPAD values under different shade conditions. Frontiers in Plant Science, 13, 928953. 10.3389/fpls.2022.928953

58. Zamani, A. S., Anand, L., Rane, K. P., Prabhu, P., Buttar, A. M., Pallathadka, H., Raghuvanshi, A., & Dugbakie, B. N. (2022). Performance of machine learning and image processing in plant leaf disease detection. Journal of Food Quality, 2022, 1598796. 10.1155/2022/1598796

59. Zhao, M. H., Li, X., Zhang, X. X., Zhang, H., & Zhao, X. Y. (2020). Mutation mechanism of leaf color in plants: A review. Forests, 11, 851. 10.3390/F11080851

