## Supplementary material for "Assessing nutritional pigment content and plant health status of green and red leafy vegetables by image analysis: Catching the “red herring” of plant digital color processing"

**Table S1** Equations of linear and non-linear regression models for estimation of chlorophyll (Chl) content using individual color features

| Color feature | Model type | Equation |
|---|---|---|
| R | Linear | Chl = -0.0118R + 2.173 |
| | Polynomial | Chl = 0.000032$(R)^2$ - 0.0186R + 2.494 |
| | Logarithmic | Chl = -1.121ln(R) + 6.1 |
| G | Linear | Chl = -0.0086G + 1.979 |
| | Polynomial | Chl = -0.000079$(G)^2$ - 0.0069G + 1.3 |
| | Logarithmic | Chl = -0.7ln(G) + 4.32 |
| B | Linear | Chl = 0.0029B + 1.04 |
| | Polynomial | Chl = -0.00041$(B)^2$ + 0.0335B + 0.55 |
| | Logarithmic | Chl = 0.15ln(B) + 0.627 |
| H | Linear | Chl = -0.0028H + 1.26 |
| | Polynomial | Chl = 0.0012$(H)^2$ - 0.093H + 2.63 |
| | Logarithmic | Chl = -0.166ln(H) + 1.75 |
| S | Linear | Chl = -0.0098S + 1.768 |
| | Polynomial | Chl = 0.000021$(S)^2$ - 0.0122S + 1.83 |
| | Logarithmic | Chl = -0.515ln(S) + 3.25 |
| I | Linear | Chl = -0.017I + 2.38 |
| | Polynomial | Chl = -0.000015$(I)^2$ - 0.0147I + 2.29 |
| | Logarithmic | Chl = -1.24ln(I) + 6.4 |
| *L** | Linear | Chl = -0.024*L** + 2.081 |
| | Polynomial | Chl = -0.00043$(L^*)^2$ + 0.011*L** + 1.46 |
| | Logarithmic | Chl = -0.816ln(*L**) + 4.089 |
| *a** | Linear | Chl = 0.0075*a** + 1.235 |
| | Polynomial | Chl = 0.000084$(a^*)^2$ + 0.0092*a** + 1.23 |
| | Logarithmic | Chl = 0.047ln(*a**) + 1.29 |
| *b** | Linear | Chl = -0.0153*b** + 1.646 |
| | Polynomial | Chl = -0.00032$(b^*)^2$ + 0.0051*b** + 1.42 |
| | Logarithmic | Chl = -0.312ln(*b**) + 2.167 |

**Table S2** Equations of linear and non-linear regression models for estimation of anthocyanin (Anth) content using individual color features

| Color feature | Model type | Equation |
|---|---|---|
| R | Linear | Anth = -0.0063R + 0.89 |
| | Polynomial | Anth = 0.000086(R)$^2$ - 0.0246R + 1.76 |
| | Logarithmic | Anth = -0.656ln(R) + 3.24 |
| G | Linear | Anth = -0.011G + 1.44 |
| | Polynomial | Anth = 0.00016(G)$^2$ - 0.044G + 2.87 |
| | Logarithmic | Anth = - 1.07ln(G) + 5.186 |
| B | Linear | Anth = 0.0255B - 0.47 |
| | Polynomial | Anth = -0.00032(B)$^2$ + 0.049B - 0.854 |
| | Logarithmic | Anth = 0.788ln(B) - 2.33 |
| H | Linear | Anth = -0.032H + 1.76 |
| | Polynomial | Anth = 0.00082(H)$^2$ - 0.095H + 2.73 |
| | Logarithmic | Anth = -1.098ln(H) + 4.4 |
| S | Linear | Anth = -0.022S + 1.757 |
| | Polynomial | Anth = 0.00044(S)$^2$ - 0.074S + 3.093 |
| | Logarithmic | Anth = -1.2ln(S) + 5.28 |
| I | Linear | Anth = -0.0126I + 1.252 |
| | Polynomial | Anth = 0.00025(I)$^2$ - 0.053I + 2.77 |
| | Logarithmic | Anth = -1.015ln(I) + 4.65 |
| *L** | Linear | Anth = -0.0276*L** + 1.42 |
| | Polynomial | Anth = 0.00098(*L**)2 - 0.107*L** + 2.867 |
| | Logarithmic | Anth = -1.076ln(*L**) + 4.23 |
| *a** | Linear | Anth = 0.042*a** + 0.9 |
| | Polynomial | Anth = 0.002(*a**)$^2$ + 0.084*a** + 0.84 |
| | Logarithmic | Anth = 0.38ln(*a**) + 0.83 |
| *b** | Linear | Anth = -0.024*b** + 1.15 |
| | Polynomial | Anth = 0.00085(*b**)$^2$ - 0.078*b** + 1.77 |
| | Logarithmic | Anth = -0.63ln(*b**) + 2.42 |
